# Pathway to Validate Gene Function in Key Bioenergy Crop, *Sorghum bicolor*

**DOI:** 10.1101/2020.12.08.416347

**Authors:** K Aregawi, J Shen, G Pierroz, C Bucheli, M Sharma, J Dahlberg, J Owiti, PG Lemaux

## Abstract

Determining gene function is an essential goal for the key bioenergy crop, *Sorghum bicolor* (L.) Moench - particularly for genes associated with its notable abiotic stress tolerances. However, detailed molecular understanding of the genes associated with those traits is limited. This was made clear in our in-depth transcriptome studies in sorghum, which indicated nearly 50% of its transcriptome is not annotated. In this report, we describe a full spectrum of tools needed to transform sorghum in order to validate and annotate genes. Efforts began with modifying a transformation method that uses the morphogenic genes *Baby Boom* and *Wuschel2* (*Ovule Development Protein2*) to accelerate transformation speed and expand amenable genotypes. In our experience, transforming RTx430 without morphogenic genes requires ~18 to 21 weeks, compared with ~10 to 12 weeks to generate T_0_ plants using methods with morphogenic genes. Utilizing morphogenic genes also allowed for the transformation of several sorghum genotypes not previously transformed or historically recalcitrant to transformation, i.e., rapid cycling SC187, stay-green BTx642, BTx623 and sweet sorghum Ramada. In order to validate candidate genes via engineering, while simultaneously introducing the morphogenic genes, a co-transformation strategy, termed altruistic transformation, was developed. To accomplish editing of the target gene, phytoene desaturase, novel constructs were created that also included morphogenic genes. To enable full characterization of transformed plants, we adapted techniques to determine copy number and independence of events at high-throughput levels. Through these efforts, we created a complete pathway from Agrobacterium infection to high-throughput molecular genotyping that can be used to ascertain gene function and expedite basic genetic research in this widely-grown bioenergy crop plant.

## Introduction

Historically, classical breeding was used to create plants with desirable new traits. However, to approach this goal more directly, modern biology has turned to engineering and editing plant genes to determine their function and ultimately to create plants with improved traits. This approach was accelerated in the model plant, *Arabidopsis thaliana*, using a simple, rapid floral-dip transformation method, which significantly facilitated the pathway to understanding gene function (Rosso et al. 2003). Later, the relative ease of engineering the model monocot, rice (*Oryza sativa* L.), also led to easier methods of determining gene function in this important crop (Wing, Purugganan, and Zhang 2018).

For many crops, such facile transformation methods (hereafter used to refer to both engineering and editing) are not available. An example is *Sorghum bicolor* (L.) Moench (sorghum), a cereal crop renowned for its drought-, heat-, and flood-tolerance. This important crop, used for food, feed, forage, and bioenergy, faces increased demand in the U.S. (TERRA 2015). One reason for its appeal as a bioenergy crop is that it can be used in all proposed renewable fuel-processing pathways: sugar, pyrolysis, cellulosic/lignocellulosic, and as a source of grain ethanol. To adequately serve bioenergy feedstock and forage needs, the function of its genes - especially those involved in its abiotic stress tolerance - need to be studied and their impact maximized. Although tolerance traits exist in specific sorghum varieties, the function of most genes controlling them is not known. Thus, it is not possible to instill optimal combinations of various abiotic stress tolerance genes into bioenergy varieties, while maintaining high biomass production.

Recent availability of large sets of transcriptomic and metabolomic data for sorghum grown under drought-stress conditions in the field (e.g., (Xu et al. 2018; Varoquaux et al. 2019; “Phytozome v13” n.d.) highlights the need for faster, more efficient methods to study gene function. Consider that although these transcriptomic data revealed that 44% of expressed genes were significantly affected by drought, a remarkable 43% of sorghum’s transcriptome is not annotated (Varoquaux et al. 2019). As such, much is left to be learned about the function of genes in sorghum’s genome.

Transient gene expression can provide insights into gene function and success in this approach has recently been achieved in sorghum (Sharma et al. 2020). However, given the complications of this approach (Ueki et al. 2013), especially under field-relevant conditions, other methods are needed to fully understand gene function in the context of the whole plant. And, even once gene function is determined, it is a lengthy and difficult process to bring together all genes needed to create an optimal bioenergy variety with abiotic stress tolerances. Thus, engineering and editing approaches are being employed to study gene function and ultimately use that information to perform marker-assisted selection or to directly introduce genes to impart desirable traits.

Performing transformation to engineer or edit sorghum is challenging because it is difficult to introduce genes into a totipotent cell that ultimately gives rise to fertile plants. Since the first plant transformation success in tobacco in 1983 (Fraley et al. 1983; Herrera-Estrella et al. 1983; Bevan, Flavell, and Chilton 1983) resulting in plants stably expressing introduced bacterial genes [for review, (Biotechnology and Biological Sciences Research Council 2013)], much progress has been made in crop plants. However, challenges remain (Altpeter et al. 2016), especially for monocots like cereals. Relative to dicots, like tobacco and Arabidopsis, monocots have been more recalcitrant to Agrobacterium-mediated transformation, largely because the natural host range of this pathogenic bacterium is dicot plants (De Cleene and De Ley 1976).

Because Agrobacterium-mediated transformation of most monocots was extremely difficult, early success in transformation of maize, for example, was accomplished using the biolistic method (Gordon-Kamm et al. 1990). Barriers to Agrobacterium-mediated monocot transformation were eventually overcome and totipotent plant cells were transformed and regenerated into fertile transformed plants. Rice was the first major monocot to be successfully transformed in this manner (Raineri et al. 1990). Additional factors, e.g., explant type, *A. tumefaciens* strain, selection methods, appropriate culture media, and amenable host genotypes, also slowed transformation success with many monocot species (Hiei, Ishida, and Komari 2014). Another issue slowing classical transformation methods was the length of time needed for tissue culture and selection, which also resulted in elevated levels of somaclonal variation, another impediment to generating improved crop species (Bregitzer, Halbert, and Lemaux 1998; Lemaux et al. 1999).

Some progress was made in addressing these difficulties in corn (Songstad et al. 1996; Frame et al. 2002; Shrawat and Lörz 2006), and rice (Ramesh, Murugiah, and Gupta 2009), however, sorghum transformation, until recently, continued to be slow and genotype-dependent. This made mutant rescue experiments or introduction of improvements into widely-grown sorghum cultivars difficult to impossible to achieve. Recent advances in maize transformation, using specific morphogenic genes under strict expression control, resulted in faster transformation methods and success with more genotypes (Lowe et al. 2016). Transformation constructs for this approach included the morphogenic genes, *Baby Boom* (*Bbm, Opd2*) and *Wuschel2* (*Wus2*). *Bbm* is an AP2/ERF transcription factor that promotes cell proliferation during embryogenesis (Boutilier et al. 2002)*. Wus2* is a homeodomain-containing transcription factor needed for maintenance of a pluripotent reservoir of stem cells in the shoot meristem (Laux et al. 1996). Overexpression of these genes in different maize cell types, under specific regulatory control, overcame in most cases genotype dependence since shoots were induced following introduction of the morphogenic genes in cells not previously able to regenerate plants (Lowe et al. 2016).

Despite the benefits of morphogenic genes being able to induce somatic embryogenesis in previously non-regenerable cell types, problems arise if those genes are not removed from transformed cells given that their continued over-expression results in negative phenotypic and reproductive outcomes in regenerated plants. To eliminate this, either excision of *Wus2* and *Bbm* or silencing of their expression is necessary after inducing embryogenesis. In early maize efforts, this was accomplished by embedding the morphogenic genes between *loxP* sites and including in the same construct CRE recombinase, driven by either the Rab17 abiotic stress-inducible promoter (Vilardell et al. 1991; Pegoraro et al. 2011), or the developmentally triggered promoter, GLB1 (Belanger and Kriz 1989). Once expressed, CRE facilitated recombination at the *loxP* sites (Sauer and Henderson 1988), causing excision of the morphogenic genes. Other successful methods of removing the morphogenic genes involved use of timing-or tissue-specific promoters to control *Wus2* and *Bbm* expression (Lowe et al. 2018). Another strategy to regenerate plants without the introduced morphogenic genes is to do co-transformation using two *Agrobacterium* strains, one with a target gene construct and the other containing the construct with the morphogenic gene(s). In this strategy, termed altruistic, most regenerated maize plants lacked integration of the morphogenic genes (Hoerster et al. 2020).

Using morphogenic genes led to successful transformation of numerous previously recalcitrant maize varieties, B73, PH2RT and HC69 (Mookkan et al. 2017; Lowe et al. 2018) and several African sorghum varieties (P. Che et al. 2018). The present report shows that use of *Bbm* and *Wus2* expands the transformable U.S. sorghum varieties, including a stay-green variety, BTx642, (Rosenow et al. 2002), the rapid cycling genotype, SC187 (Rosenow et al. 1997), the first variety with a sequenced genome, BTx623 (Frederiksen and Miller 1972), and the recalcitrant sweet sorghum Ramada (Freeman 1974). While transformation per se was successful with these genotypes, using the same construct to perform co-transformation of a second exemplary gene of interest, i.e., an RFP marker gene, was problematic. This was because *Wus2* in the construct used, pPHP81814, is driven by the maize auxin-inducible promoter 1 (*Zm-Axig1*), which does not lead to the strong expression of *Wus2* needed for co-transformation to be successful. Achieving transformation of RFP, while effectively eliminating morphogenic genes, necessitated using a different morphogenic construct. This co-transformation approach, termed altruistic in this study, required introducing into cells the altruistic morphogenic construct simultaneously with a different RFP gene-of-interest construct, in a v:v ratio of 1:9, respectively. Expression of morphogenic genes in the altruistic approach allowed those gene products to serve as facilitators, inducing the developmental pathway in cells transformed only with RFP.

After achieving success with this co-transformation approach, the morphogenic genes were included in a construct already containing the large CRISPR/Cas9 gene editing cassette to test editing efficacy. The non-altruistic approach was performed by including another exemplary gene of interest, *phytoene desaturase* (*pds*). By demonstrating editing of *pds*, this confirmed the feasibility of incorporating morphogenic genes in a very large plasmid, as well as the ability of Bbm and Wus to enhance success of CRISPR/Cas9-mediated gene editing. Taken together, the advancements described in this study provide straightforward pathways for determining gene function in sorghum using engineering and editing transformation approaches.

## Materials and Methods

### Plant materials

Seeds from sorghum varieties RTx430 (Miller 1984), BTx623 (Frederiksen and Miller 1972), BTx642 (Rosenow et al. 2002), SC187 (Rosenow et al. 1997) and the sweet sorghum Ramada (Freeman 1974) were obtained from GRIN. Seeds were planted in 3 gallon pots using SuperSoil 1 mix (Rod McClellan Co., South San Francisco, CA) and grown in the greenhouse at 28 °C with a 16h light 8h dark photoperiod. Sorghum panicles were collected at 12-14 days post-anthesis to isolate immature embryos (IEs).

### Preparation of Agrobacterium

LBA4404 Thy-is an auxotrophic (THY-) version of *A. tumefaciens* strain LBA4404 (Anand et al. 2017) into which the helper plasmid, pPHP71539 (Anand, A., Bass, S.H., Cho, H.-J., Klein, T.M., Lassner, M., McBridge, K.E. 2017), was introduced. In preparation for transformation with other constructs, streaked plates were prepared on YEP agar medium (per L: 10 g yeast extract. 10 g Bacto Peptone. 5 g NaCl, 15g Bacto-Agar, pH 7.0, 100mg/L thymidine, 50mg/L gentamicin) from bacterial stocks stored at −80 °C at 25% final concentration glycerol. After incubating in the dark for 3 d at 28 °C, 3-5 colonies from each streak plate were used to make overnight lawn cultures on YEP. Suspension cultures, prepared from lawn cultures, were diluted to OD_550_ 0.7, using PHI-I medium (Wu et al. 2014) with 0.005 % silwet and 0.2 mM acetosyringone.

### Transformation constructs

All primers used for constructs in this section are listed in **Supplementary Table 1.** The pPHP81814 construct (Chu et al. 2019) contains the maize morphogenic genes *Wuschel2* (*Wus2*) (Nardmann and Werr 2006) and *Baby Boom* (*Bbm*) (Boutilier et al. 2002), as well as the screenable marker gene ZSGreen1 (*ZSG*), driven by the maize auxin-inducible promoter, *Zm-Axig1*, the maize phospholipid transfer protein promoter, *Zm-PLTP*, and the *S. bicolor* Ubiquitin promoter, *Sb-UBI*, respectively. These three genes are contained within Loxp sites; *cre* recombinase is driven by the maize late embryogenesis-stage GLB1 promoter (Belanger and Kriz 1989). The acetolactate synthase (*ALS*) selectable marker gene is driven by the *S. bicolor* acetolactate synthase promoter. The pPHP71539 “helper” plasmid contains *virB*, *virC*, *virD*, *virE* and *virG* operons and a bacterial gentamicin resistance gene (Anand et al. 2018).

To create the altruistic construct, pGL190 (**Figure 1A**), pPHP83911 (**Supplementary Figure 1A**) was the entry vector, containing the sorghum Ubi promoter, *uidA* and sorghum gamma kafirin terminator, which was linearized between attL3 and attL4, using pPL01 and pPL04 primers. The expression cassette, including the sorghum Ubi1 promoter and its first intron, ZSG, and the rice Ubi terminator, was then inserted using pPL02 and pPL03 primers via Gibson assembly. The entry vector was verified using Sanger sequencing with primers pPL02 and pPL05, and M13 reverse primers. To complete pGL190 construction, the entry vector was recombined with destination vector, pPHP85425 (**Supplementary Figure 1B**), and confirmed by *BamHI* restriction. pGL190 was electroporated into LBA4404 Thy-; plasmid from positive colonies was confirmed by complete plasmid sequencing (Center for Computational and Integrative Biology, Massachusetts General Hospital, Boston MA).

**Figure 1.**
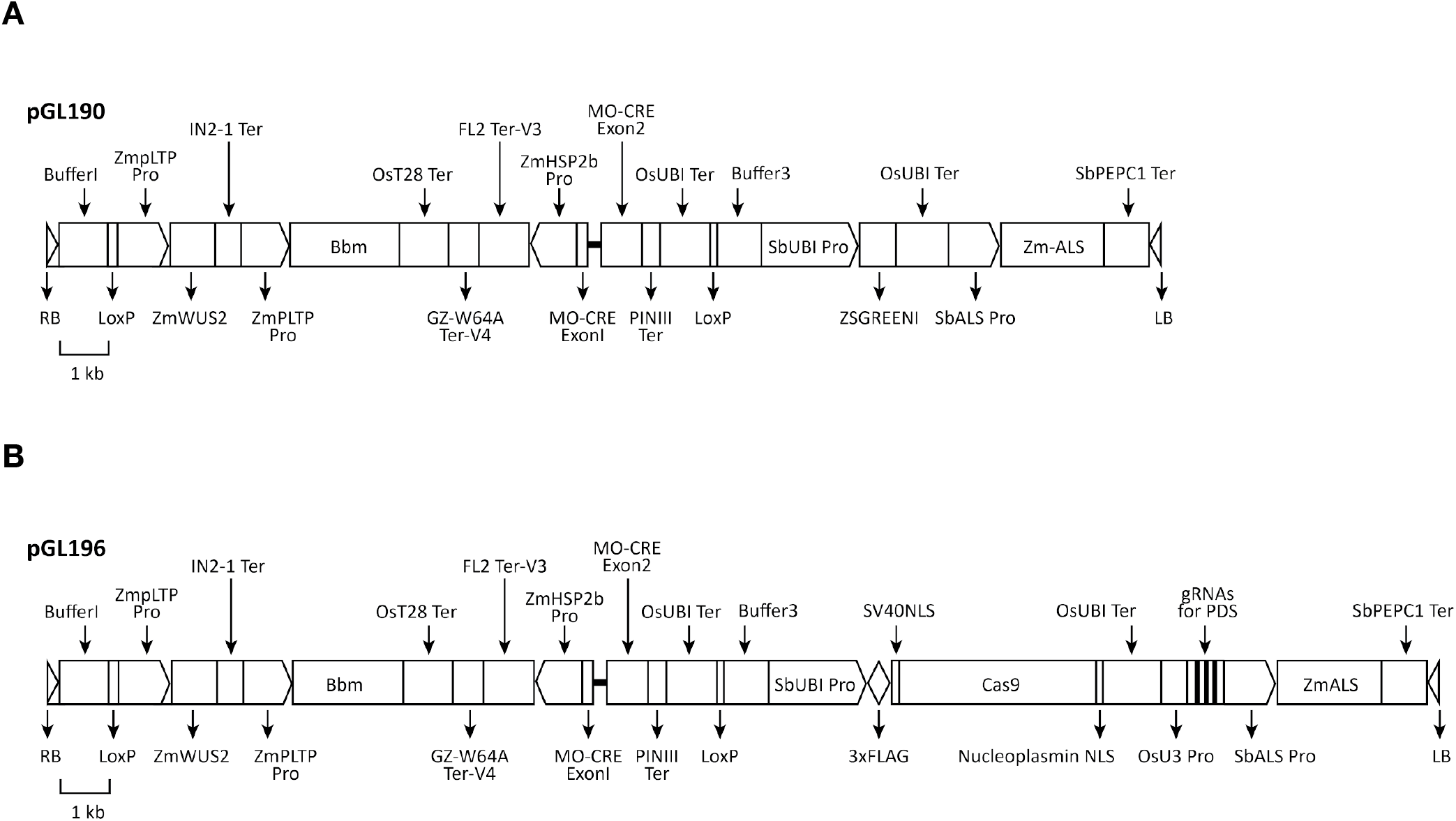
Construct maps for transformation. **A.** pGL190 for altruistic engineering. **B.** pGL196 for non-altruistic editing of phytoene desaturase genes. Triangles indicate T-DNA borders. Rectangles with arrows indicate promoters. Blank rectangles indicate genes or buffer regions. Black rectangles indicate terminator regions.

For pGL193 (**Supplementary Figure 1C**), pPHP83911 (**Supplementary Figure 1A**) was amplified with pPL14 and pPL15 primers; Cas9 was amplified from pRGEB32 with pPL12 and pPL13 primers. These two fragments were ligated via Gibson assembly (New England BioLabs, Ipswich MA). The gRNA expression unit, amplified from pRGEB32 with pPL16 and pPL17 primers, was inserted via Gibson assembly into the *Sma*I-linearized Gibson assembly vector. Three tRNA guide RNAs (gRNAs), targeting phytoene desaturase gene exons in sorghum (**Figure 2A**), were amplified from pGTR (Xie, Minkenberg, and Yang 2015) with pPL18 to pPL23. The tRNA-gRNA units were generated using Golden Gate assembly, digested by *Fok*I, and ligated into pGL193 for the LR recombination reaction. pGL196 (**Figure 1B**) was generated by recombining the *pds* tRNA-gRNA units and Cas9 with pPHP85425 (**Supplementary Figure 1B**). For pGL198, the maize ubiquitin promoter and maize codon-optimized SpCas9 gene were amplified from pBUN411 (Xing et al. 2014) with the primers pPL31 and pPL32, and the sorghum ubiquitin promoter and SpCas9 gene on pGL193 were replaced by Gibson assembly (**Supplementary Figure 1D**). All constructs were confirmed by Sanger sequencing. The destination vectors, pGL190 (**Figure 1A**) and pGL196 (**Figure 1B**), isolated from Agrobacterium, were confirmed by complete plasmid sequencing (Center for Computational and Integrative Biology, Massachusetts General Hospital, Boston MA).

**Figure 2.**
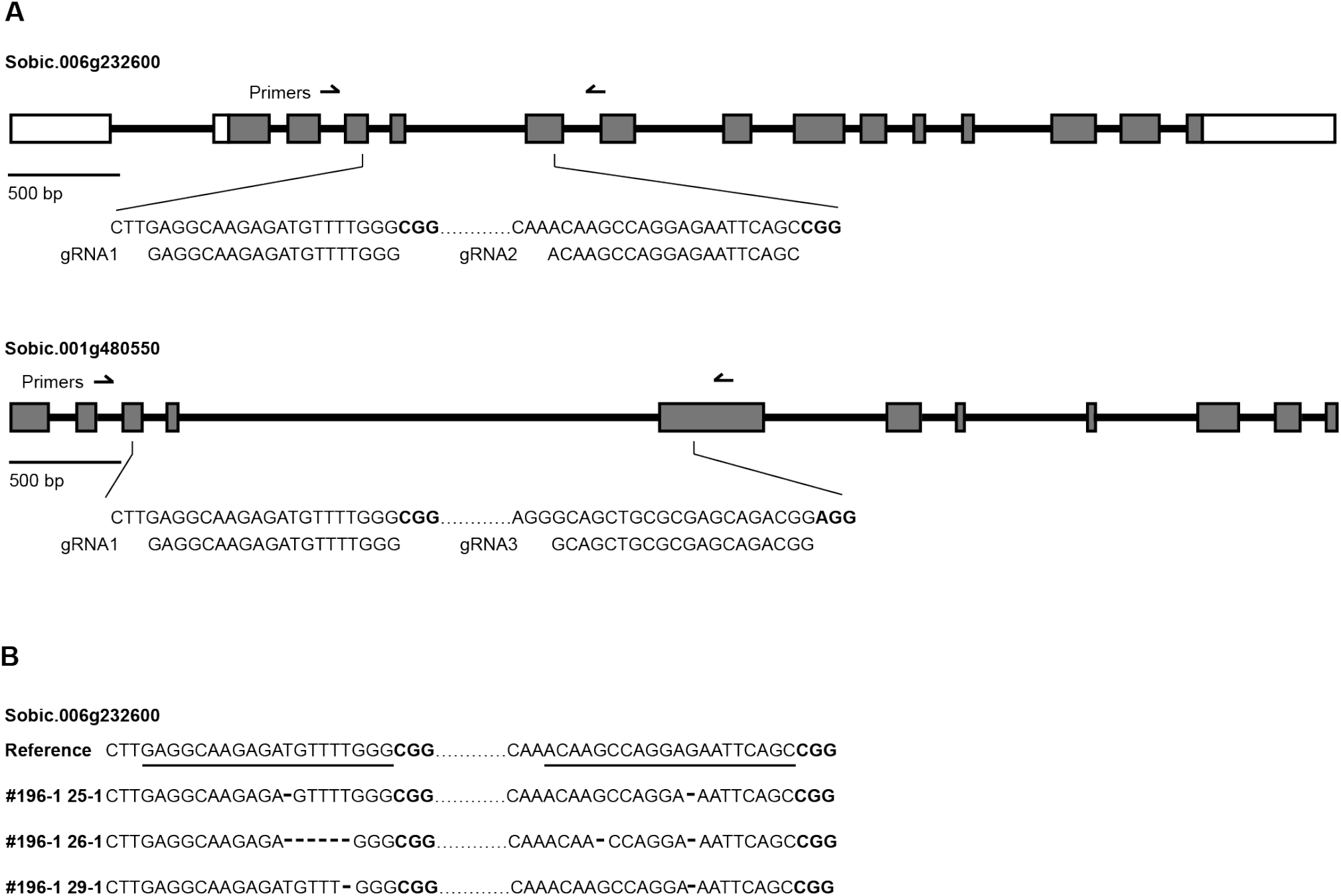
Editing of *pds* genes using CRISPR/Cas9 system. **A.** Three guide RNAs (gRNAs) targeting *pds* genes. White rectangles indicate untranslated regions; grey rectangles indicate exons; black lines indicate intron regions. Arrows indicate primers for genotyping. gRNA1 is located in the third exons of pds genes. gRNA2 and gRNA3 are located in the fifth exons of pds genes. PAM sites for gRNAs are shown in bold black font. **B.** Sanger sequencing of edited pds genes. gRNAs are underlined and PAM sites for gRNAs are shown in bold black font. Deletions are shown with dashes.

### Non-altruistic *A. tumefaciens*-mediated sorghum transformation

All media used in this experiment, modified from (Jones et al. 2019), are described in **Supplementary Table 2**. A general description of constructs, plant and bacterial selection genes and their purpose is in **Supplementary Table 3.** Immature seeds of RTx430, BTx623, BTx642, SC187 and Ramada were isolated from greenhouse-grown plants, surface-sterilized twice for two mins with 75% ethanol, then 20% bleach plus 0.2% Tween20 for twenty mins. IEs of 1.5-2 mm were isolated and put into PHI-I liquid medium. Once all IEs were isolated, PHI-I was removed, 1ml of Agrobacterium suspensions was added and mixed for 5 min on a shaker at medium speed and then the suspension was removed by pipetting. IEs were placed on co-cultivation medium (CCM) scutellum side up, and kept for 7 d at 24 °C in the dark. IEs were transferred to resting medium for 7 d to halt Agro growth; tissues were then moved to embryo maturation media (EMM) with 0.05 mg/L imazapyr (IMZ; Sigma Aldrich Chemicals, St. Louis MO) until shoots formed (**Supplementary Figures 2A,B**). Plantlets (2-3 cm) were moved to rooting medium (RM) with 0.05 mg/L IMZ under 16-h photoperiod at 26 °C (**Supplementary Figure 2C**). Plants with well-established roots and shoots (10-12 cm in height) were moved to soil in the growth chamber at 28 °C with 16-h light and 8-h dark photoperiod for conditioning for two weeks before transferring to the greenhouse. Percent transformation efficiency was calculated as the total number of T_0_ plants that are PCR-positive for the target gene, divided by the total number of IEs used times 100.

### Altruistic *A. tumefaciens*-mediated sorghum transformation

Immature seeds of RTx430 were isolated and sterilized; IEs were excised as described for non-altruistic transformation. For transformation, Agrobacterium LBA4404 Thy-, containing either pGL190 or pANIC10A at OD_550_ 0.7, were mixed in a 1:9 (pGL190:pANIC10A) volume:volume ratio. IEs were cultured as described for non-altruistic transformation except, rather than selecting with IMZ as for non-altruistic transformation, hygromycin (hyg; PhytoTechnology Laboratories, Lenexa KS) at 20mg/L, was added to EMM; no selection agent was used during rooting. Brightfield and fluorescence microscopy images were taken. pGL190 and pANIC10A alone were also used as controls for transformation.

### DNA extraction

Leaf samples, ~3 cm, were frozen in liquid nitrogen and ground in a MM300 bead beater (Retsch GmbH, Haan Germany) for 1.5 mins at 25 cps. Samples were then re-frozen and ground again; 500 ul of urea buffer (2M urea, 0.35mM NaCl, 20mM Tris-HCl pH 8, 20mM EDTA, 1% sarkosyl) was added and tubes vortexed for 30 secs. 10ul of RNase A was added, incubated 10 mins at room temperature and 500 ul of phenol:chloroform:IAA (25:24:1) was added, vortexed 15 mins, and centrifuged 13,800 x g for 15 mins. 550 ul of the aqueous phase was transferred to a new tube and 55 ul of 3M sodium acetate (pH 5.2) and 367 ul of isopropanol were added; tubes were inverted to precipitate DNA and incubated at −20 °C overnight or −80 °C for 1 h. After centrifugation at 13,800 x g for 5 mins, supernatant was removed and pellet rinsed in 500 ul 70% ETOH. Samples were centrifuged for 5 mins at 13,800 x g and supernatant removed; samples were centrifuged a final time for 1 min and excess ethanol removed. Pellets were air dried 5-10 mins in laminar flow hood until clear. Pellets were resuspended in sterile distilled water and stored at −20 °C.

### Fluorescence visualization

Zeiss Lumar v12 epifluorescence stereomicroscope and QImaging Retiga SRV camera were used to observe IEs expressing *RFP* and *ZSG* at 3 to 10 d post-Agrobacterium infection, while tissues were on EMM. RFP and ZSG fluorescences were detected using Texas Red and Endow GFP filter sets, respectively.

### PCR analysis of putative transgenic events

PCR was performed using genomic DNA isolated from leaf tissue of T_0_ putative transformed plants. For pPHP81814 putative, non-altruistic transformants, PCR primers (**Supplementary Table 4**) were used to confirm presence of *ZSG* and *ALS*, which is reinstated following CRE-mediated excision of *BBM* and *WUS*. For altruistic transformation with pGL190 and pANIC10A, PCR was performed as described using *hph* and *Bbm* primers (**Supplementary Tables 4, 5**).

### Transgene copy number analysis

To determine transgene copy number, primers and HEX™-labeled probes were designed to detect the endogenous reference gene, protein phosphatase 2A (PP2A) (**Supplementary Table 4**). The PP2A reference gene, chosen as most stable in *Sorghum bicolor* under different stress treatments and tissues (Sudhakar Reddy et al. 2016), was used for analysis of RTx430, SC187 and Ramada. A different primer set (**Supplementary Table 6**) was used to detect acetolactate synthase (ALS), using FAM™-labeled probes and double-quenched with ZEN™ and Iowa Black Hole Quencher^®^ (Integrated DNA Technologies, Inc., Coralville, IA). Probes and primers were designed according to Bio-Rad Laboratories criteria; droplets were generated using the QX200 droplet generator (Bio-Rad Laboratories, Hercules CA). Samples included no template control (ddH20), negative control (wild-type sorghum genomic DNA) and a positive control (plasmid DNA), in addition to genomic DNA from transformed plants. HindIII, which does not cut the transgene, was used to digest genomic DNA. Samples were transferred to 96-well plates; a thermal cycler was used for PCR amplification. To optimize quantification of transgene copy number by digital droplet PCR, gradient PCR (55 °C - 65 °C) and dilution gradient of sorghum DNA (6.8 ng - 55 ng) was used to determine optimal annealing temperatures and DNA amounts. Among those annealing temperatures and DNA amounts, 55 °C and 27.5 ng genomic DNA were used because they gave the highest fluorescence amplitude difference between negative and positive droplets, specific amplification and clear separation between negative and positive droplets. PCR started with 95 °C enzyme activation for 10 mins, followed by 94 °C for 30 secs and 55 °C for 1 min; each step was performed with 40 cycles and 2 °C/second ramp rate. The process was finalized by enzyme deactivation at 98 °C for 10 mins. Following PCR, fluorescence data was measured using QX200 Droplet reader (Bio-Rad Laboratories). More than 10,000 droplets were generated for each sample. Samples were performed in duplicate; data was analyzed by QuantaSoft™ Software (Bio-Rad Laboratories).

### Independent event determination

All primers used in this section are listed in **Supplementary Table 1.** The method for determining independence of events was modified from (O’Malley et al. 2007) as follows. Adaptors for *Hin*dIII (adpH3) and *Bam*HI (adpB1) were annealed with pPL24 and pPL25 and pPL24 and pPL26, respectively. Oligonucleotides were heated to 98 °C in a water bath then cooled to room temperature by turning the water bath off. 1 ug genomic DNA was added to adpH3 and adpB1 adapters, using *Hin*dIII and *Bam*HI and T4 ligase; the reaction was carried out at 25 °C for 16 h in a thermocycler. After ligation, PCR amplification was performed using pPL27 and pPL28 primers at 60 °C for annealing; elongation was for 4 min. If distinct bands were not seen, PCR products were used as templates for nested-PCR amplification as follows. 1 ul of PCR product was amplified at 60 °C for annealing, elongation for 4 min, using pPL29 and pPL30 primers. PCR products, purified with QIAquick PCR purification kit (QIAGEN, Germantown MD), were confirmed by Sanger sequencing with LBb1 as the sequencing primer. Independent insertion location was determined using BLAST (NCBI Resource Coordinators 2016). In modifying from (O’Malley et al. 2007) to determine transgene independence, genomic DNA amounts were increased from the 30 ng used for Arabidopsis to 900 ng for sorghum because of its larger genome size; digestion/ligation reaction temperatures remain at 25 °C. Also, PCR amplification for T-DNA and genomic DNA junctions were optimized individually to obtain high success rates. For the process of determining independence of events to be successful, it is not mandatory to first verify plants are single-copy; however, this makes amplification and sequencing easier.

### gRNAs for CRISPR-Cas9 gene editing

To demonstrate editing efficacy, three gRNAs targeting *phytoene desaturase,* Sobic.006G232600, on chromosome 6, and its closest homolog, Sobic.001G480550, on chromosome 1 were generated (**Figures 2A,B**). The first gRNA targeted the third exon of both genes; the second and third gRNAs targeted the fifth exon. For use in non-altruistic editing, gRNAs were ligated with transfer RNAs in pGL193 (**Supplementary Figure 1B**).

For the non-altruistic editing method that includes the use of morphogenic genes, a Cas9 expression cassette was used in which Cas9 expression is driven by the sorghum ubiquitin promoter and gRNA-tRNA expression is driven by rice snoRNA U3 promoter, which is on the pENTR vector, pGL193 (**Supplementary Figure 1B**). After incorporating gRNAs targeting *pds* (**Figures 2A,B**), both cassettes were introduced into the destination vector, pHP85425 (**Supplementary Figure 1D)**, yielding, pGL196 (**Figure 1B)**. pGL196 was transformed into sorghum cultivar RTx430, employing Agrobacterium strain LBA4404 Thy-. Right before moving tissues to EMM, IEs on resting media were treated with heat shock at 45 °C and 75% humidity for 2 hrs to trigger morphogenic gene excision. From each IE, 3-4 plantlets were moved into rooting medium and PCR-tested; only one was sequenced and moved to soil.

## Results

### Non-altruistic *A. tumefaciens*-mediated transformation

Using the non-altruistic transformation method, five sorghum genotypes: RTx430, SC187, BTx642, BTx623 and Ramada were successfully transformed. At early stages of the transformation process, insertion and expression was confirmed by visualizing in IEs expression of the fluorescence ZsGreen (ZSG) marker gene, using brightfield and fluorescence microscopy at 3-10 days post-transformation while tissues were on EMM (**Figures 3A,B**). Many somatic embryos strongly expressed ZSG (**Figure 3B**). After being transferred to rooting medium, putative transformed plants were genotyped using PCR. While transformed plants were obtained from all genotypes used, efficiencies varied widely among the different genotypes (**Table 1**). Of the five genotypes, the highest efficiency of T_0_ plants positive only for the ALS gene was obtained using a standard sorghum transformation variety RTx430 (9.4%), followed by the previously non-transformed rapid-cycling variety, SC187 (2.0%). Also transformed were the stay-green variety, BTx642 (0.9%) and BTx623 (0.1%), the first sorghum genotype with a published complete genome sequence (Paterson et al. 2009). One of the most previously recalcitrant varieties, sweet sorghum Ramada, has a published transformation efficiency of 0.09% (Raghuwanshi and Birch 2010). In the present study an efficiency of 1.2% was achieved, an approximate 10-fold improvement. It is also noteworthy that the time required to generate RTx430 T0 plants, for example, was reduced in this approach to ~10-12 weeks, compared to the typical 18-21 weeks reported in (Wu et al. 2014) and also observed in unpublished work in our laboratory. For the non-altruistic transformation method, imazapyr (IMZ), a nonselective, systemic herbicide targeting acetolactate synthase, was used as the selection agent in both EMM and rooting media. Most IEs going through IMZ selection had at least some tissue for both RTx430 (**Figures 3C,D,E**) and BTx642 that remained healthy and was not browning or necrotic (**Supplementary Figures 2A,B**). However, once on rooting medium most untransformed plantlets turned brown and died as a result of selection. Although the use of IMZ resulted in effective selection, more escapes were observed relative to the hygromycin selection used for altruistic transformation (see Altruistic Transformation section). Additionally, RTx430 and BTx642 responded differently to selection, especially on EMM. For instance, RTx430 tissues which were not killed by selection on EMM formed more somatic embryos (**Figures 3D,E**) and continuously proliferated to form large-sized tissue pieces, about 1 cm in diameter. However, BTx642 tissues, relative to RTx430, formed a smaller number of somatic embryos and did not proliferate to be as large as RTx430 tissues (**Supplementary Figures 2A,B**). Notably, IMZ selection had a stronger effect on tissues of BTx642 than on those of RTx430.

**Table 1.**
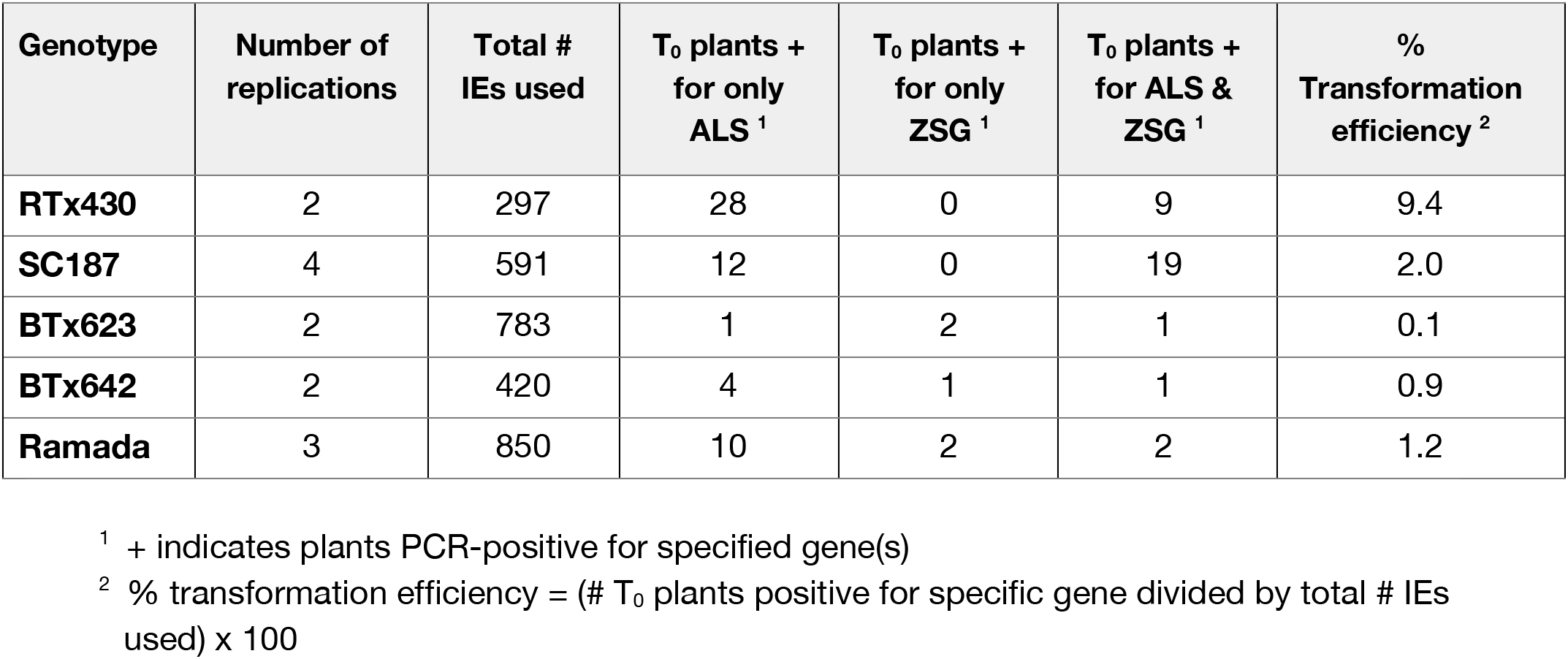
Non-altruistic transformation efficiency of five sorghum genotypes using pPHP81814 and IMZ selection.

**Figure 3.**
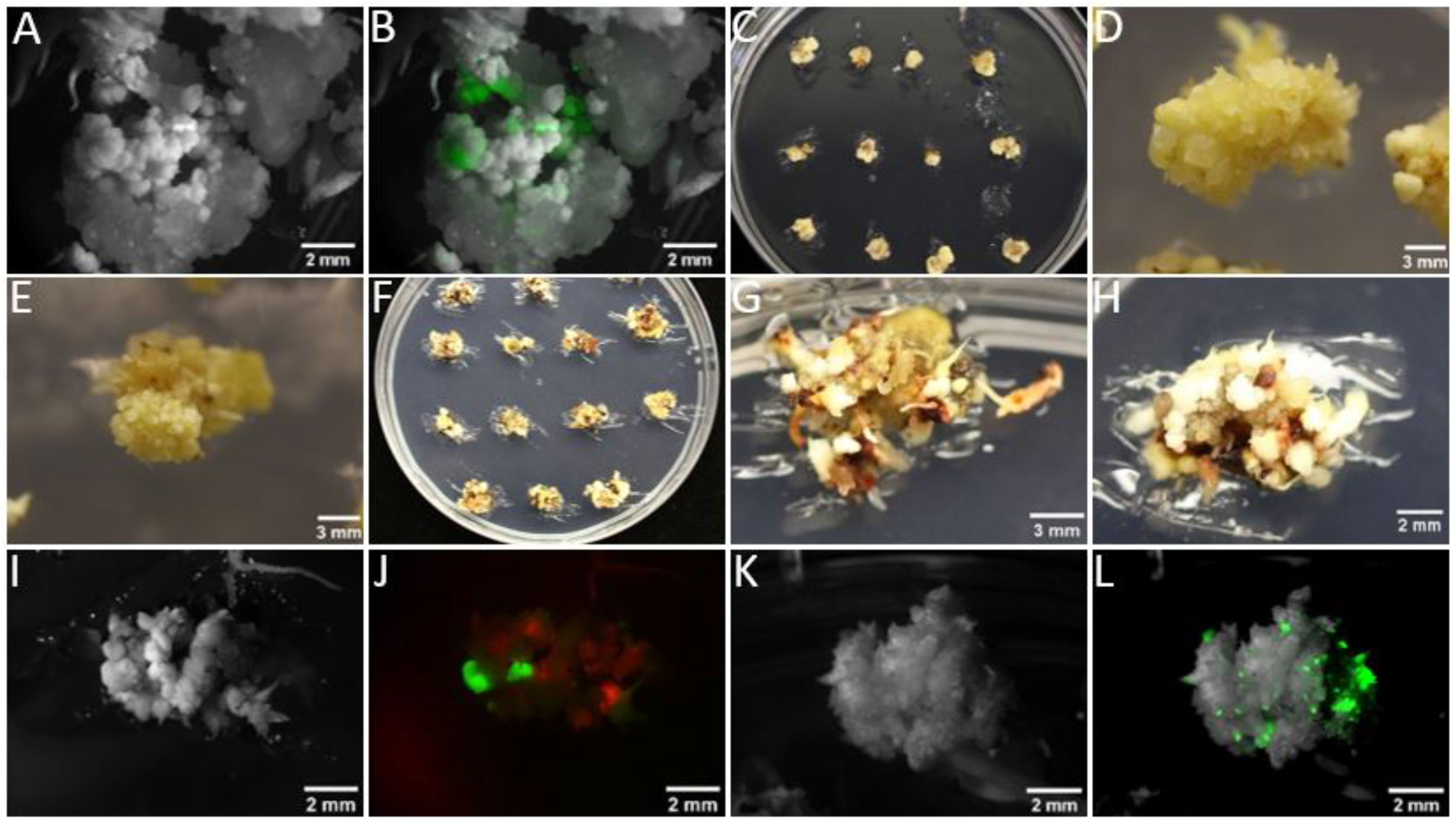
RTx430 IEs, 1.5-2mm, were transformed with Agrobacterium LBA4404 Thy-strain, containing appropriate constructs depending on the experiment. IEs were cultured for one week each on co-cultivation then resting medium. For non-altruistic transformation, using pPHP81814 (ZSG), images by **A.** brightfield and by **B.**fluorescence were taken while tissues were on EMM; in **B.** ZSG expression was observed. **C.D.E.** For non-altruistic transformation RTx430 tissues were moved to EMM with 0.05 mg/L imazapyr (IMZ). **F.G.H.** For altruistic transformation 20mg/L hygromycin was used as a selection agent for RTx430. Images were taken using **I.K.** brightfield and **J.L.** fluorescence microscopy. **J.** For altruistic transformation, using pGL190 (ZSG) and pANIC10A (RFP), IEs expressed ZSG and RFP. **I.** For comparison a brightfield image of the same tissue is shown. As a control, pGL190 alone was used to transform RTx430 IEs and imaged using **K.** Brightfield and **L.** fluorescence; ZSG expression in transformed tissue is shown.

### Genotypes and phenotypes of T_0_ plants and T_1_ seed set

While inclusion of morphogenic genes in transformation constructs is necessary to achieve transformation success with more sorghum genotypes, those genes must be removed from transformed tissues to achieve plant regeneration and normal phenotypes and seed set. For non-altruistic transformation with pPHP81814, putative transformed plants were tested by PCR, using appropriate primers (**Supplementary Table 4**), to confirm removal of ZSG, *Wus2* and *Bbm* by CRE-mediated excision, achievable because all three genes are flanked by *loxP* sites. Different genotypes showed different percentages of morphogenic gene excision (**Table 1**). Highest excision rates were for Ramada, 83.3%, BTx642, 80.0% and RTx430, 75.5%. For BTx623 and SC187 rates were 50.0% and 38.7%, respectively. Additionally, because portions of introduced morphogenic genes might remain, visual assessment of T_0_ plants at late development was performed to see if any phenotypic abnormalities were observed. Plants negative for introduced morphogenic genes showed normal growth and seed set (**Supplementary Figures 3A,B**). As anticipated, plants testing positive for introduced morphogenic genes showed abnormal phenotypes, e.g., short stature, twisted leaves and/or poor seed set; similar results for maize were previously reported (Lowe et al. 2016). Identical observations were made for plants resulting from altruistic transformation.

### Transgene copy number analysis

Digital droplet PCR (ddPCR) was used to quantify transgene copy numbers. Values generated for copy number were close to integer values and results revealed that the majority of transformed plants were single-copy: 63.2% of RTx430 (12 of 19), 75.0% of SC187 (9 of 12), and 77.8% (7 of 9) of Ramada plants (**Table 2**). The remainder of the plants appeared to be multi-copy.

**Table 2.**
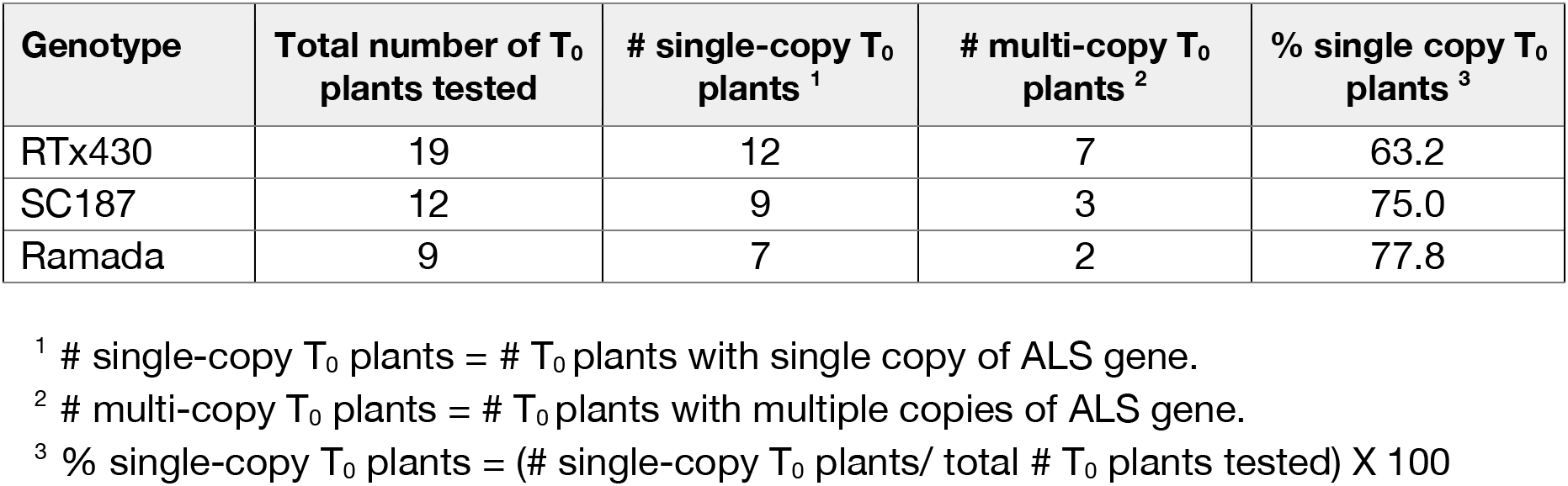
Transgene copy number in T_0_ plants of different sorghum genotypes determined using digital droplet PCR.

### Determining independence of transformed events

The adaptor ligation-mediated PCR method, used for mapping T-DNA inserts in Arabidopsis (O’Malley et al. 2007), was modified to determine independence of different transformed plants. To test the method, six transformed plants from the same IE and two transformed plants from different IEs were chosen. PCR product bands were all of the same size for the plants from the same IE, indicating that they were not independent events (**Figure 4A**), likely resulting from tillering. However, bands from transformed plants from different IEs were of a different size, showing that they were independent events, as expected. Thus, although transformed plants arising from the same IE can derive from tissue having independent insertions, these results suggest that they also might not be independent, since multiple shoots can arise from different original cells in the same somatic embryo as it proliferates. Sequencing results provided additional information on mapping, showing that the insertion from one line was in one gene, Sobic.006G073700, while the insertion in another event was in an intergenic region (**Figure 4B**).

**Figure 4.**
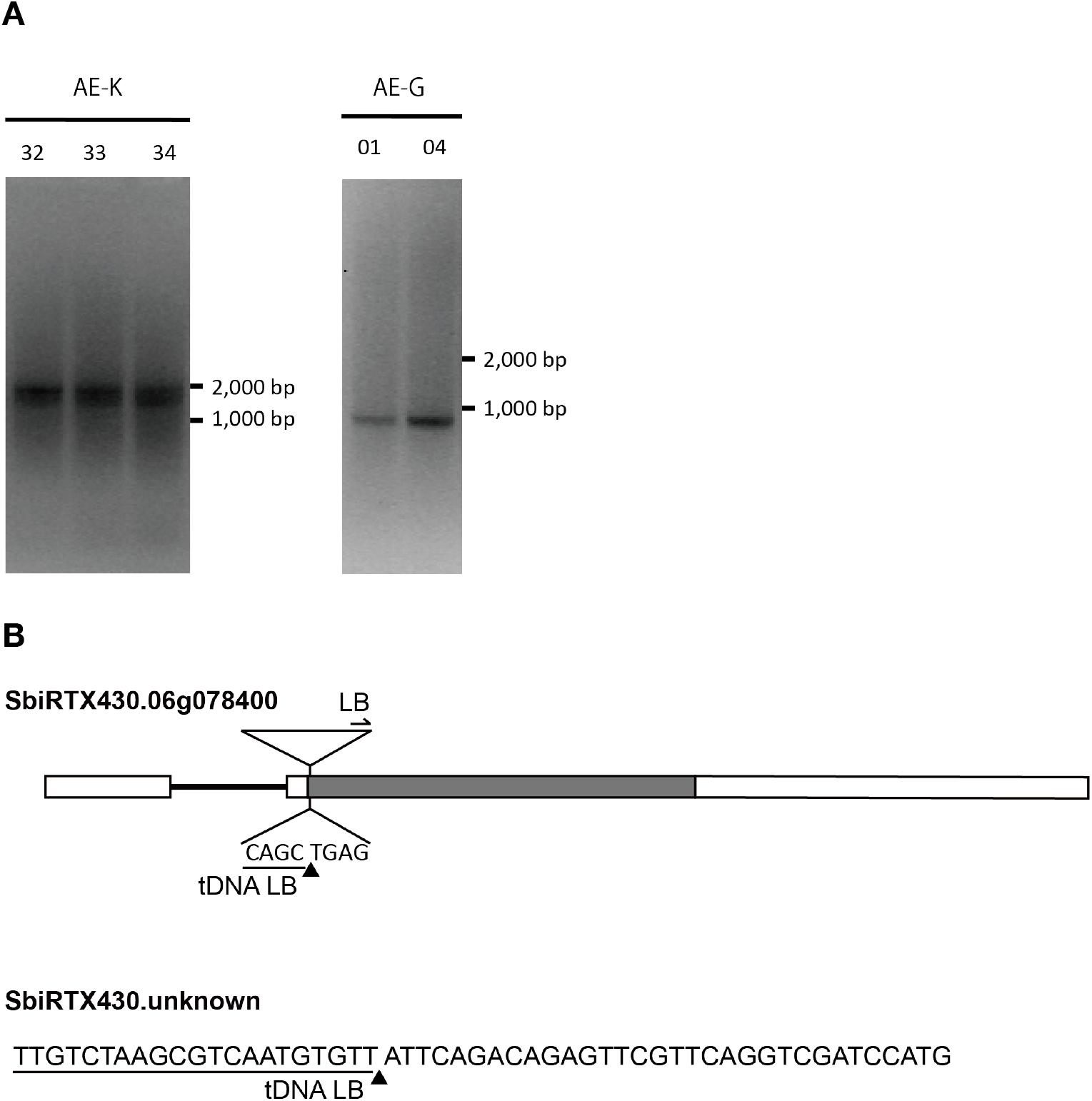
Independent event determination in altruistic engineering events. **A.** Gel images used to determine independent event status. Genomic DNA and T-DNA conjunctions were amplified by PCR. DNA ladder locations are shown on the right. **B.** Examples of T-DNA insertion sites on the sorghum genome. T-DNA insertions were confirmed by Sanger sequencing. Arrow in upper image indicates left border primer. Triangles indicate T-DNA insertion sites. T-DNA left border sequences are underlined.

### Altruistic *A. tumefaciens-*mediated transformation

RTx430 IEs were transformed using modifications of the altruistic method described for maize (Hoerster et al. 2020). The altruistic approach required sorghum IEs to be transformed simultaneously with two Agrobacterium strains. One contained our “gene of interest”, RFP, and a selection gene, hygromycin phosphotransferase (*hph*) in the pANIC10A plasmid (Mann et al. 2012). The second Agrobacterium strain contained pGL190, which had the morphogenic genes, *Wus2* and *Bbm*, and ZSG. The two Agro strains were mixed 9:1 v:v, pANIC10A:pGL190. The altruistic approach required use of a different selection scheme from that used for the non-altruistic approach, which was based on IMZ. For the altruistic effort, tissues were selected on EMM containing hygromycin (**Figures 3F,G,H**), to select for expression of *hph* in pANIC10A, which also had the RFP gene; no selection was used on rooting media. Unlike IMZ selection, hygromycin selection was stronger, having visible effects on the tissues, i.e., growth of non-transformed tissues was strongly inhibited and became necrotic. The strong selection effect of hygromycin was also apparent in the genotyping results, given that, even though no selection agent was included in rooting media, there was only a small number of escapes in contrast to results from IMZ selection. To follow expression of ZSG from pGL190, which also contained the morphogenic genes, and RFP from pANIC10A, both brightfield (**Figures 3I**) and fluorescence microscopy images (**Figures 3J**) were used. Transformation with only pGL190 with ZSG expression (**Figures 3K,L**) and pANIC10A were used as controls.

It was important to determine efficiencies of the altruistic approach, both for transformation in general and for editing, and these different applications need to be considered separately. This is prudent because different outcomes could benefit more from one approach compared to the other. Thus, if a true independent event was the goal, as when doing classical engineering to overexpress a gene, choosing a single event from each IE would provide an effective way to assure independently transformed plants. If multiple independent edited plants is the goal, maximizing the number of independent events would not be the means to generate the largest number of independent edits. Randomly choosing plants and then genotyping to identify unique edits would likely yield larger numbers of novel edits, since editing can occur at random times during the growth of transformed tissue.

For the first method, aimed at generating multiple independent events, two transformation replications were conducted using 95 and 84 IEs (**Table 3A**). Transformation efficiencies in the replications were 23.2% and 28.6%, respectively. The adaptor ligation-mediated PCR method, used for determining independence of events, will be used to show whether plants from the same IE were the same or independent. Results from previous experiments suggest that individual plants from the same IE may be independent or not, if they arose via branching or tillering.

**Table 3.** Altruistic transformation efficiency for RTx430 using pGL190 and pANIC10A

**A**
| Replication | Total # IEs used | T<sub>0</sub> plants + for only RFP <sup>1</sup> | T<sub>0</sub> plants + for only Bbm <sup>1</sup> | T<sub>0</sub> plants + for hph & Bbm <sup>1</sup> | Transformation efficiency (%) <sup>2</sup> |
| --- | --- | --- | --- | --- | --- |
| 1 | 95 | 22 | 0 | 7 | 23.2 |
| 2 | 84 | 24 | 4 | 9 | 28.6 |

<sup>1</sup> + indicates plants PCR-positive for specified gene(s)
<sup>2</sup> Transformation efficiency (%) = (# T<sub>0</sub> plants + for only RFP divided by total # of IEs used) x 100

**B**
| Replication | Total # IEs used | T<sub>0</sub> plants + for only hph <sup>1</sup> | T<sub>0</sub> plants + for only Bbm <sup>1</sup> | T<sub>0</sub> plants + for hph & Bbm <sup>1</sup> | Transformation efficiency (%) <sup>2</sup> |
| --- | --- | --- | --- | --- | --- |
| 1 | 100 | 52 | 13 | 25 | 52.0 |
| 2 | 100 | 15 | 1 | 10 | 15.0 |

<sup>1</sup> + indicates plants PCR-positive for specified gene(s)
<sup>2</sup> Transformation efficiency (%) = (# T<sub>0</sub> plants + for only RFP divided by total # of IEs used) x 100

In the second approach, which would be used if the aim were to maximize the number of edits, two replications were performed using 100 IEs each. For this method, any plantlet of sufficient size was transferred from EMM and then to rooting media, regardless of whether another plantlet had been recovered previously from that same IE. This approach is appropriate for an editing experiment, since editing is not limited to a particular point in time but can happen at any time during tissue proliferation. In this approach, the focus would be on first determining if a plant is transformed using PCR for the introduced gene(s) and then positive plants would be genotyped and sequenced to identify independent edits.

In this scenario, for the second approach, 134 individual plantlets, following selection with hygromycin, were regenerated from two replications. Of those randomly selected plantlets, 67 were positive only for *hph*, which is linked to the “gene of interest” RFP; 14 were positive only for *Bbm*; 35 were positive for both *hph* and *Bbm* (**Table 3B**). In this scenario, the transformation efficiency, which reflects a plant containing only the “gene of interest” and not that it was independent, was 33.5% (67/200). At this point if an editing construct was used, as described in the next section, those positive plants would be sequenced to determine novel editing efficiency. Using this approach should maximize editing efficiency.

### CRISPR-Cas9 editing

Success rates of methods used for CRISPR/Cas9-mediated editing of *pds*, employing Cas9 and gRNAs, were determined using a single non-altruistic construct with morphogenic genes and the entire CRISPR/Cas9 editing cassette, pGL196. (**Figure 1B**). This construct was introduced into RTx430, and transformed tissue was heat-treated to trigger excision of morphogenic genes. From two replications, 95/76 IEs were used and 17/12 plants tested positive for only gRNAs (**Table 4**). Based on the number of IEs used, this yielded 17.9% (17/95) to 15.8% (12/76) transformation efficiencies, respectively. Of those plants positive only for gRNAs from the two reps, six and three were edited, yielding 35.3% (6/17) or 25.0% (3/12) of gRNA+ plants being edited. As a percent of the total number of IEs used, editing efficiencies were 6.3% (6/95) to 3.9% (3/76). To analyze editing events, the *pds* sequence on chromosome 6 was determined and it was found that both gRNA targeting sites were edited in three events; three were only edited on gRNA1 **(Figure 2B).** The albino phenotype, expected following biallelic editing, was not observed in T_0_ plants, so those plants were moved to soil and T_1_ plants will be analyzed for albino phenotypes, to confirm functional *pds* knockouts. Since editing frequency was not high enough to efficiently generate edited plants and biallelic edited plants were not observed, the Cas9 expression cassette was modified to include a maize ubiquitin promoter driving maize codon-optimized Cas9 to improve editing efficiency (**Supplementary Figure 1D**) and is currently being used to transform RTx430 to determine efficiencies of biallelic knockouts.

**Table 4.** Non-altruistic editing efficiency for RTx430 using pGL196

| Replication | Total # IEs used | T<sub>0</sub> plants + for gRNAs only <sup>1</sup> | T<sub>0</sub> plants + for gRNA & Bbm <sup>1</sup> | % Transformation efficiency - gRNA only <sup>2</sup> | # Edited plants | % Editing efficiency per IE <sup>3</sup> | % Editing efficiency per gRNA + transformant <sup>4</sup> |
| --- | --- | --- | --- | --- | --- | --- | --- |
| 1 | 95 | 17 | 14 | 17.9 | 6 | 6.3 | 35.3 |
| 2 | 76 | 12 | 15 | 15.8 | 3 | 3.9 | 25.0 |

<sup>1</sup> + indicates T<sub>0</sub> plants PCR-positive for specified gene(s)
<sup>2</sup> % Transformation efficiency = (# T<sub>0</sub> plants +ve for only gRNA divided by total # of IEs used) x 100
<sup>3</sup> % Editing efficiency = (# edited plants divided by total # of IEs used) x 100
<sup>4</sup> % Editing efficiency = (# edited plants divided by total # of T<sub>0</sub> plants + for gRNAs only) x 100

## Discussion

To accelerate gene function studies in important crop plants, improvements in transformation are needed (Altpeter et al. 2016). This became obvious to us when our transcriptome data analysis of drought-stressed field-grown sorghum revealed that, although 44% of expressed genes were significantly affected by drought, 43% of sorghum’s transcriptome is not annotated (Varoquaux et al. 2019). One pathway to annotating these genes is determining their function using genetic engineering and editing.

Among the major impediments to performing such genetic modification of sorghum, when using classical transformation methods without morphogenic genes, are genotype-dependence and the long time periods needed for selection, regeneration, and obtaining seed from the transformants. In the present study, we overcame some of these challenges. First, we used constructs containing the morphogenic genes, *Bbm* and *Wus2*, and in initial studies we used a non-altruistic transformation method that led to T_0_ transformed sorghum plants 8-9 weeks earlier than when the construct did not contain morphogenic genes [Lemaux et al., unpublished; (Wu et al. 2014)]. Also, use of morphogenic genes for transformation increased the number of sorghum varieties amenable to transformation. This advance made it possible to use a fast-cycling variety, SC187, from which T1 plants can be generated ten weeks faster than when using the standard transformation variety, RTx430. That rapid generation time is also seen in field data with average days to booting for RTx430 being 93.4d (s.d. 2.4) versus 49.8d (s.d. 2.6) for SC187 (F. Harmon, personal communication). Availability of a high-quality Pac-Bio sequence for SC187 (“Phytozome v13” n.d.) facilitates gene editing efforts in this variety. In sum, these advances provide distinct advantages over earlier transformation methods for gene function studies.

Successfully transforming different sorghum genotypes by introducing morphogenic genes that are expressed at specific levels and under specific conditions led to formation of somatic embryos directly from surface cells of the scutellum of an IE (Lowe et al. 2018). This obviated an extended callus phase, a situation which can reduce somaclonal variation. Additionally, using this method expanded the number of sorghum varieties amenable to transformation, facilitating, for example, complementation studies of mutants present in previously non-transformable varieties. The present study reports the first successful transformation of BTx642, the prototype stay-green sorghum variety, providing opportunities for gene function studies related to this important trait. Transformation of sweet sorghum Ramada, at a rate approximately 10-times the frequency previously published (Raghuwanshi and Birch 2010), will enable determining the function of genes involved in stem sugar production. High quality PacBio reference sequences for BTx642, SC187 and RTx430 (“Phytozome v13” n.d.), along with effective transformation strategies, will provide increased capacity to determine the molecular bases of certain important traits.

While development of such methods has been critical in advancing the capacity to transform monocot species, increasingly such efforts are aimed at determining the function of uncharacterized genes. An apparent challenge when using the non-altruistic transformation method to study gene function was the inability to identify transformed plants with the gene of interest but devoid of introduced morphogenic genes that cause developmental abnormalities (**Supplementary Figure 3**). A recent study described an altruistic transformation method aimed at avoiding this outcome (Hoerster et al. 2020). In this approach high expression levels of *Wus2,* but not *Bbm,* in cells in close proximity to those with the introduced gene of interest yielded plants with the gene of interest but not the morphogenic genes. Success of this method was described as depending on transformed cells diffusing enough *Wus*2 into neighboring cells containing only the gene of interest in order to trigger developmental progression. In this situation the neighboring cell with the gene of interest can divide and ultimately regenerate a transformed plant with the gene of interest but without having introduced morphogenic genes.

For the altruistic strategy to be successful, strength of the promoter driving the morphogenic gene is critical. In successful non-altruistic transformation efforts in maize, the phospholipid transferase protein gene (*Pltp*) promoter, which supports strong expression in the scutellar epithelial layer of IEs (Lowe et al. 2018), was used to drive *Bbm* expression, but not *Wus2*. Alternatively, for successful altruistic transformation of maize, the *Pltp* promoter was used to drive *Wus2* while *Bbm* was not expressed (Hoerster et al. 2020). In the present study, for the non-altruistic method using pPHP81814, the promoter driving *Wus2 expression* was the maize auxin-inducible promoter, AXIG1 promoter (Garnaat, Lowe, and Roth 2002; Lowe et al. 2018). This promoter does not support sufficiently strong expression levels to trigger developmental progression in neighboring cells (unpublished results). Thus, for successful altruistic engineering in sorghum, constructs were created in which the *Pltp* promoter drove expression of both *Wus2* and *Bbm*. In these efforts a 9:1 ratio of Agrobacterium carrying the exemplary gene of interest in pANIC10A, RFP, compared to the construct with the morphogenic genes was used. This ratio was shown in maize to increase likelihood of identifying transformed plants with only the gene of interest (Hoerster et al. 2020). With the present approach to altruistic transformation, it was necessary to optimize selection of tissue with hygromycin, not IMZ that was used in our non-altruistic approach, because *hph* was linked to the RFP gene of interest.

For altruistic engineering, using pGL190 and pANIC10A, transformed RFP plants were generated without morphogenic genes; however, efficiencies varied widely between replicates (**Tables 3A,B**). In the maize effort viral enhancers were used in the *pltp::wus2* cassette to increase Wus2 expression in transformed cells to such a level that presence of the Wus2 protein itself likely hindered regeneration of those plantlets with only the morphogenic gene (Hoerster et al. 2020). Given that large numbers of plants with both pGL190 and pANIC10A (**Tables 3A,B**) were identified in the present study, it is likely that Wus2 expression was not strong enough to be toxic to transformed cells, leading to survival of plants with morphogenic genes with or without RFP. Regardless, transformation speed and efficiency was enhanced greatly by using the co-transformation technique. So, this strategy can be utilized to increase speed of transformation by employing previously completed gene-of-interest plasmids without needing to insert morphogenic genes into the same construct.

From previous experiments (data not shown) and pGL196 editing experiments (**Figure 1B**), it was observed that the non-altruistic morphogenic strategy led to tissues forming more somatic embryos, which proliferated and gave plantlets faster than with the altruistic strategy. Furthermore, because altruistic engineering results were inconsistent, the feasibility of combining morphogenic genes with another large expression cassette, specifically the CRISPR/Cas9 editing cassette, was tested. The resulting non-altruistic editing strategy also has disadvantages. Because the non-altruistic construct includes morphogenic genes as well as the entire CRISPR/Cas9 editing cassette, the resulting plasmid is very large, ~36kb. So, synthesis of similarly sized plasmids, as well as their introduction into Agrobacterium, can prove difficult. The non-altruistic strategy also necessitates removal of introduced morphogenic genes using heat shock treatment, which compared to the altruistic approach leads to a higher ratio of regenerated plants positive for both morphogenic genes and the gene of interest relative to plants positive solely for the gene of interest (**Tables 3,4**).

For transformed plants to be created effectively for gene function studies, copy number and independence of events must be established. In the present study digital droplet PCR was used to determine copy number and in the three genotypes tested, over 60% of independent transformed plants were single-copy. Determining whether events are independent is also particularly important with sorghum because of its tendency to tiller, making it unclear if plants from the same IE are clonal or independent. Previous methods to determine independence of events were Southern hybridization and Southern-by-Sequencing (Zastrow-Hayes et al. 2015), both of which can be time-and resource-intensive. We developed a high-throughput method to assess independence of events by modifying a method that was previously used for transgene insertion mapping in Arabidopsis (O’Malley et al. 2007).

Previous reports on editing in sorghum described success with endogenous genes, using both biolistic bombardment and classical Agrobacterium-mediated transformation (Che et al. 2018; Li et al. 2018; Liu, Li, and Godwin 2019; Char et al. 2020). To improve transformation and subsequent editing efficiencies in sorghum, we used a morphogenic gene-mediated non-altruistic transformation strategy, using multiple guide RNAs to edit an exemplary gene of interest, *pds,* with the CRISPR/Cas9 system. Numerous transformants were identified which did not integrate morphogenic genes; however, the expected albino phenotype, caused by a *pds* knockout, was not observed in T0 generation plants. Several reasons for this are possible. First, editing efficiency using pGL196 (**Figure 1B**) was not as high as expected, likely due to low-level expression of Cas9, driven by the sorghum ubiquitin promoter (SbUBI) (I. Godwin, personal communication). Use of pGL198 (**Supplementary Figure 1C**), in which SbUBI and SpCas9 were replaced by the ZmUBI and maize codon-optimized SpCas9 will provide a more likely path to increased editing efficiency. Additionally, the existence of chimeras upon editing leads to rare phenotypic changes in T0 plants.

Introducing a single, approximate 36 kB construct, including gRNAs, Cas9, LoxP and morphogenic genes, into sorghum was shown in this study to lead to an approximate 5% editing efficiency - providing a path for editing using one large construct with all required elements. But, there are possible ways to increase rates of generating heritable editing mutations, for example, using a meiosis-specific promoter, like the dmc1 promoter used in maize genome editing (Feng et al. 2018), to drive Cas9. Also, Cas9 orthologs or variants, like Streptococcus thermophilus Cas9 (StCas9) (Müller et al. 2016) and Staphylococcus aureus Cas9 (SaCas9) (Ran et al. 2015), can be used to improve editing frequency in sorghum.

## Conclusion

While certain steps have been demonstrated that pave the way to using engineering and editing to improve crop plants and to increasing agricultural productivity, the critical next step is consolidating those improved technologies to have a clear path from introduction of target genes to determining physiological function. As described in this study, use of morphogenic genes led to a wider pool of sorghum genotypes amenable to transformation, relative to previous efforts. Given that success, the challenge facing researchers with limited resources is being able to identify independent, single-copy events devoid of the introduced morphogenic genes. This was accomplished in the present study using existing constructs (Corteva Agriscience) and creating new morphogenic constructs. Modifications needed to achieve success included identification of successful selection and culturing methods for altruistic and non-altruistic transformation, coupled with creation of new editing constructs that are compatible with the use of morphogenic genes. We provide detailed descriptions of these resources and methods – including transformation of additional sorghum genotypes, the identification of single-copy, independent events, and the creation of non-altruistic editing constructs containing morphogenic genes. These advancements have led to a complete pathway to identify gene function in sorghum – paving the way to improved plants to meet climate and population challenges.

## Online content

Supplementary figures and tables are available in the Supplementary Materials.

## Acknowledgement

This project was supported by the Department of Energy, Biological and Environmental Research Program, Competitive Grant DE-SC0014081. M.S. acknowledges support from DBT India and IUSSTF as a B-ACER fellowship. PGL acknowledges support from the US Cooperative Extension Service through the Division of Agriculture and Natural Resources of the University of California. We thank Drs. William Gordon-Kamm and Ping Che at Corteva Agriscience for helpful conversations and for the constructs pPHP71539, pPHP81814, pPHP83911 and pPHP85425. We also thank Dr. Ronan O’Malley for helpful discussions on the use of his transgene insertion mapping technique. We are grateful for the help of the staff of the UC Berkeley Biological Imaging Facility and the UC Berkeley Oxford Tract Greenhouse Facility. We also thank Barbara Alonso for help in creating Fig. 1 and for managing the reference list.

## Author Contributions

Kiflom Aregawi: Investigation, Transformation Methodology, Writing - Original Draft - Review & Editing. Jianqiang Shen: Investigation, Transformation Methodology, Constructs, Writing - Review & Editing. Grady Pierroz: Investigation, Transformation Methodology, Writing – Review & Editing. Claudia Bucheli: Investigation, Writing – Review & Editing. Manoj Sharma: Transformation Methodology, Writing - Review & Editing. Jeffery Dahlberg: Methodology. Judith Owiti: Transformation Methodology, Writing - Review & Editing. Peggy G. Lemaux: Conceptualization, Formal analysis, Supervision, Project administration and funding acquisition, Writing – Original Draft –Review & Editing.

## Data availability statement

Accession numbers and gene names are provided, as available, in the text and **Supplementary Table 7.** Methods for generation of different constructs and transformation protocols are described in the text and in Supplementary Figures. PHP vectors are freely available for research, but commercial applications require a paid non-exclusive license from Corteva Agriscience.

