## Supplementary Figures for "Pathway to Validate Gene Function in Key Bioenergy Crop, *Sorghum bicolor*"

**A.**

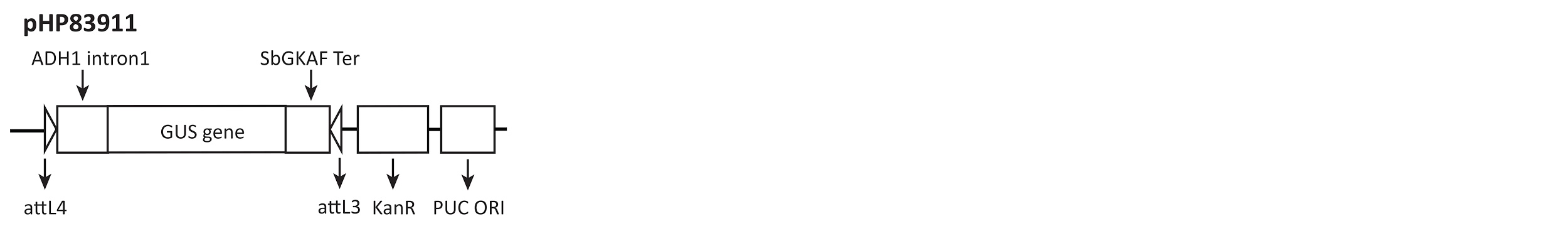

**B.**

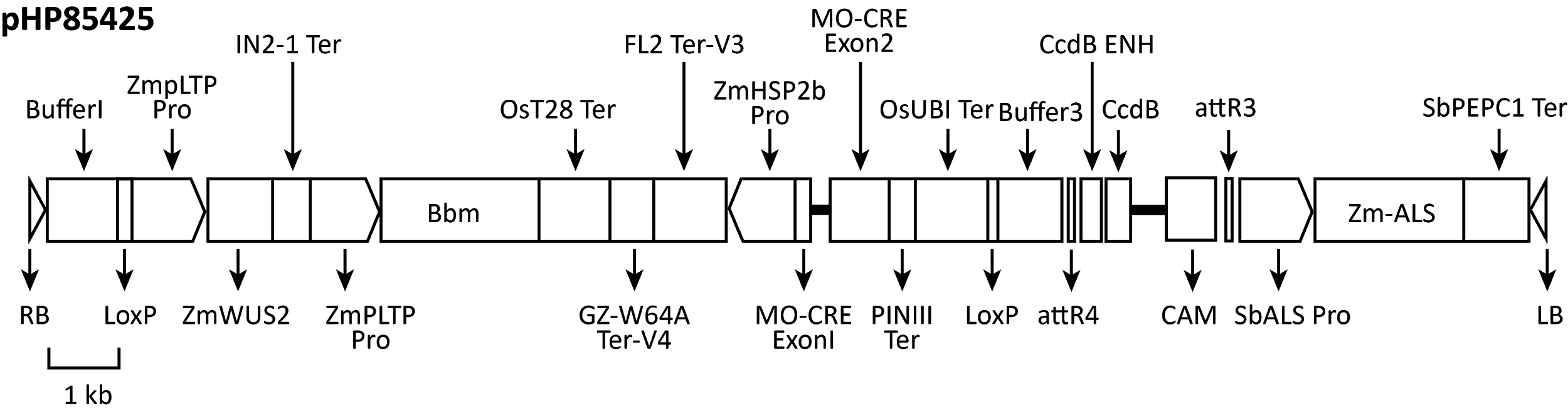

**C****.**

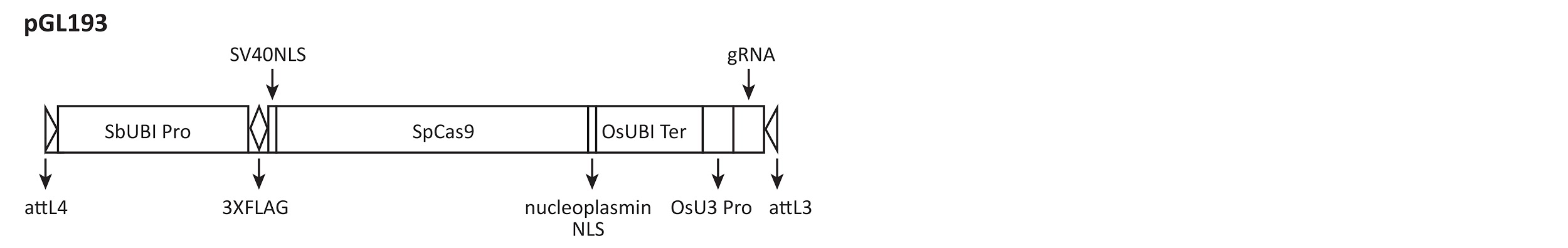

**D.**

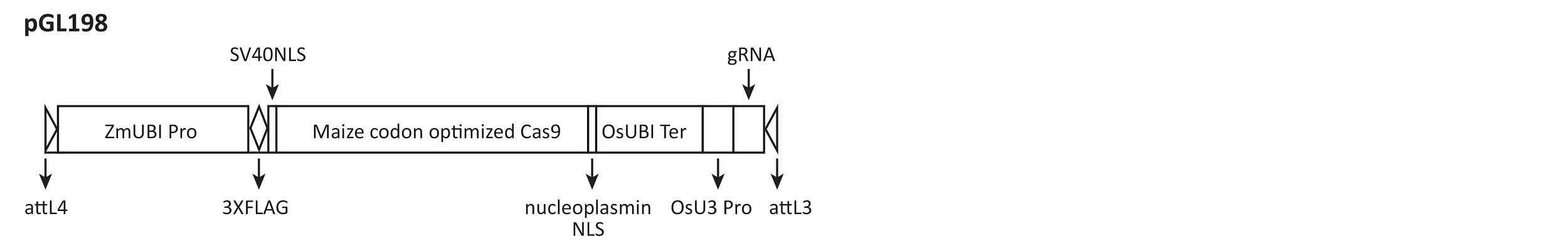

**Supplementary Figure 1. A.** pPHP83911, used for generating pGL193 and pGL198, is a pENTR vector. **B.** pPHP85425 is a destination vector for non-altruistic transformation. **C.** pGL193 is a pENTR vector for non-altruistic editing, with gRNAs. **D.** pGL198 is a pENTR vector for non-altruistic editing, with the maize ubiquitin promoter and maize codon-optimized SpCas9.

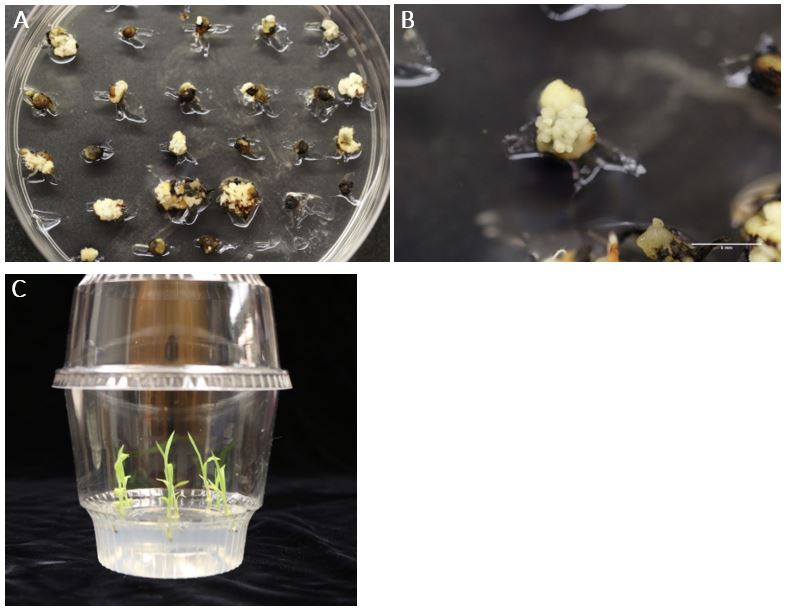

**Supplementary Figure 2. A. B.** BTx642 tissues on EMM with IMZ selection following transformation with pPHP81814. **C.** Plantlets growing on rooting media with IMZ.

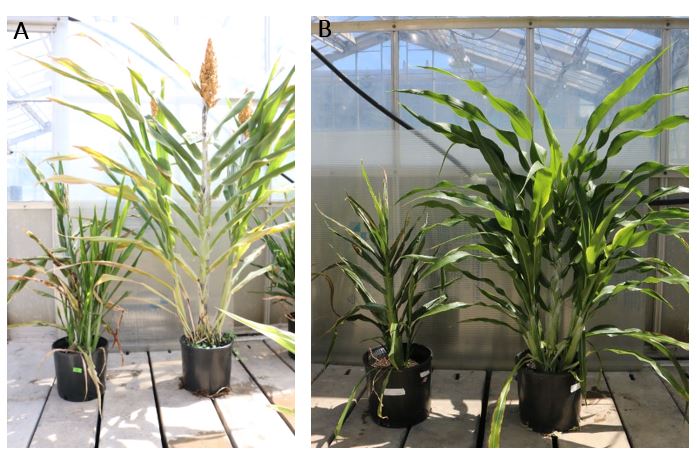

**Supplementary Figure 3.** Plants from **A.** RTx430 and **B.** BTx642 transformed with pPHP81814. In both **A., B.** wild-type plants (right) and plants retaining introduced developmental genes (left) have shorter stature, twisted leaves and poorer seed set, compared to wild-type.
