## Supplementary Tables for "Pathway to Validate Gene Function in Key Bioenergy Crop, *Sorghum bicolor*"

**Supplementary Table 1.** Primers for constructs, gRNAs, determining independent events

| **Accession**  **Name** | **Purpose** | **Primer Name** | **Sequences 5 -> 3** |
| --- | --- | --- | --- |
| pPL01 | pGL190 construction | attL4 R | caacttttctatacaaagttggcattataaaaaagcattgcttatcaat |
| pPL02 | “ | attL4_pUBI F | gccaactttgtatagaaaagttgctgcccttaaggccaattgttcaagattcattcaaca |
| pPL03 | “ | attL3_tUBI R | ggtgaccggcgcgccgaagcctaccaaagcaaagcgtttgtcga |
| pPL04 | “ | attL3 F | ggcgcgccggtcacccggt |
| pPL05 | “ | ZS Green 2 R | accctatcgagatctgagtc |
| pPL06 | pGL193 construction | Cas9 F | ctttaacttagcctaggatccgactataaggaccacgacggagactacaaggatca |
| pPL07 | “ | Cas9 R | actggccattggtaccctttttcttttttgcctggccggcctttttcgtg |
| pPL08 | “ | pENTR F | ggtaccaatggccagttaacagatccagct |
| pPL09 | “ | pENTR R | ggatcctaggctaagttaaagtcgacctgcag |
| pPL10 | “ | gRNA unit F | cctagaaggccacccagcggataacaatttcacacaggaaacagctatgaca |
| pPL11 | “ | gRNA unit R | aactttgtataataaagttgcccttgtggacctgcaggcatgcacg |
| pPL12 | gRNAs for sorghum PDS | PDS-gR1 F | taGGTCTCCAGATGTTTTGGGgttttagagctagaa ^1^ |
| pPL13 | “ | PDS-gR1 R | cgGGTCTCAATCTCTTGCCTCtgcaccagccggg ^1^ |
| pPL14 | “ | PDS-gR2 F | taGGTCTCCGGAGAATTCAGCgttttagagctagaa ^1^ |
| pPL15 | “ | PDS-gR2 R | cgGGTCTCACTCCTGGCTTGTtgcaccagccggg ^1^ |
| pPL16 | “ | PDS-gR3 F | taGGTCTCCGCGAGCAGACGGgttttagagctagaa ^1^ |
| pPL17 | “ | PDS-gR3 R | cgGGTCTCATCGCGCAGCTGCtgcaccagccggg ^1^ |
| pPL18 | Determining independent event | LSA 1 | gtaatacgactcactatagggcacgcgtggtcgacggcccgggctgc |
| pPL19 | “ | SSAH3 | phosphate-AGCTGCAGCCCG-amino C7 ^2^ |
| pPL20 | “ | SSAE1 | phosphate-AATTGCAGCCCG-amino C7 ^2^ |
| pPL21 | “ | LBa1 | tcacaattccacacaacatacgagcc |
| pPL22 | “ | AP1 | gtaatacgactcactatagggcacgcg |
| pPL23 | “ | LBb1 | attaattgcgttgcgctcactgccc |
| pPL24 | “ | AP2 | tggtcgacggcccgggctgc |
| pPL25 | pGL198 construction | ZM-UBI F | gccaactttgtatagaaaagttgctgcagtgcagcgtgacccg |
| pPL26 | “ | ZM-opt Cas9 R | actggccattggtacccttcttcttcttcgcctgccccgc |

^1^ Capitalized letters indicate gRNA sequences.

^2^ Capitalized letters indicate oligonucleotides.

#### **Supplementary** **Table 2.** Medium composition for sorghum transformation

| **Medium name** | **Medium components** |
| --- | --- |
| PHI-I (infection medium) | MS salts plus vitamins, 4.43 g/L, thiamine-HCl 1mg/l, 2,4-D 1.5 mg/l, sucrose 68.5 g/l, glucose 36 g/l, acetosyringone^1^ 39.24 mg/l, pH 5.2. |
| Co-cultivation | MS salts plus vitamins, 4.43 g/L, thiamine-HCl 1mg/l, 2,4-D 2 mg/l, sucrose 20 g/l, glucose 10 g/l, L-proline 0.7 g/l, MES buffer 0.5 g/l, acetosyringone^1^ 39.24 mg/l, ascorbic acid^1^ 10 mg/l, thymidine^1^ 100mg/l, pH 5.8. |
| Resting | MS salts plus vitamins, 4.43 g/L, thiamine-HCl 1 mg/l, 2,4-D 2 mg/l, sucrose 20 g/l, glucose 10 g/l, L-proline 0.7 g/l, MES buffer 0.5 g/l, acetosyringone^1^ 39.24 mg/l, ascorbic acid^1^ 10 mg/l, carbenicillin^1^ 25 mg/l, pH 5.8. |
| ^2^ Embryo-maturation medium (EMM) | MS salts plus vitamins, 4.43 g/L, zeatin 0.05 mg/l, copper sulfate 1.25mg/l, L-proline 0.7 g/l, sucrose 60 g/l, IAA^1^ 1 mg/l , ABA^1^ 0.026mg/l , thidiazuron 0.1 mg/l, BAP^1^ 1 mg/l, carbenicillin^1^ 250mg/l, pH 5.6. |
| ^2^ Rooting | MS salts 4.43 g/l, sucrose 40 g/l, pH 5.6 |

##### ^1^ Added after medium is autoclaved and cooled to ~55℃

##### ^2^ For EMM and rooting medium, selection agent (hyg or IMZ) will change depending on construct

#### **Supplementary Table 3.** Constructs used for sorghum transformation

| **Construct** | **Purpose** | **Plant selectable marker gene (selection agent)** | **Bacterial selectable marker gene (selection agent)** |
| --- | --- | --- | --- |
| pPHP81814 | Non-altruistic transformation | Acetolactate synthase, ALS (Imazapyr) | Adenylyltransferease, aadA (spectinomycin) |
| pANIC10A | Altruistic transformation | hygromycin phosphotransferase, hph (hygromycin) | Aminoglycoside phosphotransferase, nptIi, (kanamycin) |
| pGL190 | Altruistic transformation | Acetolactate synthase, ALS (Imazapyr) | Adenylyltransferase, aadA (spectinomycin) |
| pGL192 | Entry vector | NA | Aminoglycoside phosphotransferase, nptII, (kanamycin) |
| pGL193 | Entry vector for gRNAs targeting PDS | NA | Aminoglycoside phosphotransferase, nptII, (kanamycin) |
| pGL196 | CRISPR editing target for PDS, non-altruistic | Acetolactate synthase, ALS | Adenylyltransferase, aadA (spectinomycin) |
| pGL197 | CRISPR editing target for PDS, altruistic | Hygromycin phosphotransferase, hph, (hygromycin) | Aminoglycoside phosphotransferase, nptII, (kanamycin) |
| pGL198 | Entry vector | NA | Aminoglycoside phosphotransferase, nptII, (kanamycin) |

#### **Supplementary** **Table 4.** Primers for genotyping putative transformed plants

| **Gene symbol** | **Gene Name** | **Construct name** | **Primers (F/R) (5’-3’)** | **Amplicon length (bp)** | **Tm (^o^C)** |
| --- | --- | --- | --- | --- | --- |
| *ALS2* | Acetolactate  synthase | pPHP81814 | CTTTGGCTCATGGAACGA  ATCTTCTTTATCGCTGCGC | 576 | 58.2 |
| *ZSG* | ZsGreen | pPHP81814 | CTCCTGCGAGAAGATCATC  ACCCTATCGAGATCTGAGTC | 277 | 55 |
| *RFP* | Red fluorescence  protein | pANIC10A | GGCCATTATACGTGCGACTT  GCATGTGCAATTTCTCGTTG | 155 | 60 |
| *Bbm* | Baby boom | 190 | GCCGGAGCAACCACTACAT  TCCGCCACCATTGTTCTC | 132 | 61 |
| *hph* | Hygromycin phosphotransferase | pANIC10A | GAAGAATCTCGTGCTTTCAGCTTCG  CAAGCTCTGATAGAGTTGGTCAAGACC | 741 | 64 |
| *Bbm* | Baby boom | pGL190/  pGL196 | GGTCGTCAAGTCTATTTAGGTGGCT  AAGTAAAGATCCTTGTTCCCTGCAACT | 299 | 63 |
| *gRNA unit* | gRNA-tRNA | pGL196 | CCTAGAAGGCCACCCAGCGGATAACAATTTCACACAGGAAACAGCTATGACA  AACTTTGTATAATAAAGTTGCCCTTGTGGACCTGCAGGCATGCACG | 1,157 | 60 |
| *PDS-06g* | Phytoene desaturase | NA | TGTAAGTTGGGGAATTTCGAGGGAACCACTA  GCAGAGTTAAGCTCAACGGTAATGGTGTAGTG | 1,127 | 60 |

#### **Supplementary** **Table 5.** PCR programs for genotyping putative transformed plants

| **Gene symbol** | **Gene Name** | **Construct name** | **Initial Denaturation** | **Denaturation** | **Annealing** | **Extension** | **Extension** | **Hold** |
| --- | --- | --- | --- | --- | --- | --- | --- | --- |
| *ALS2* | Acetolactate synthase | pPHP81814 | 95^o^C-3:00min | 95^o^C-0:30sec | 54^o^C-0:30sec | 72^o^C-1:00 min | 72^o^C-5:00 min | 12^o^C- ∞ |
| *ZSG* | ZsGreen | pPHP81814 | 95^o^C-4:00min | 95o^o^C-0:30sec | 55^o^C-0:30sec | 72^o^C-0:50  sec | 72^o^C-7:00 min | 12^o^C- ∞ |
| *RFP* | Red fluorescence protein | pANIC10A | 95^o^C-3:00min | 95^o^C-0:30sec | 53^o^C-0:30sec | 72^o^C-0:30  sec | 72^o^C-5:00 min | 4^o^C- ∞ |
| *Bbm* | Baby boom | pGL190 | 95^o^C-3:00min | 95^o^C-0:30sec | 53^o^C-0:30sec | 72^o^C-0:30  sec | 72^o^C-5:00 min | 4^o^C- ∞ |
| *Hph +*  *Bbm* | Hygromycin phosphotransferase + Baby boom | pGL190 +  pANIC10A | 95^o^C-3:00min | 95^o^C-0:30sec | 57^o^C-0:30sec | 72^o^C-1:00  min | 72^o^C-5:00 min | 4^o^C- ∞ |
| *gRNA unit* | gRNA-tRNA | pGL196 | 98^o^C-3:00min | 98^o^C-0:10sec | 60^o^C-0:15sec | 68^o^C-1:15  min | 68^o^C-5:00 min | 4^o^C- ∞ |
| *PDS* | Phytoene desaturase | NA | 98^o^C-3:00min | 98^o^C-0:10sec | 60^o^C-0:15sec | 68^o^C-1:15  min | 68^o^C-5:00 min | 4^o^C- ∞ |

### **Supplementary Table 6.** Primers and probes for digital droplet PCR

| **Gene symbol** | **Gene name** | **Primers (F/R) (5’-3’)** | **Probe sequence (5’-3’)** | **Amplicon length (bp)** | **Tm (^o^C)** |
| --- | --- | --- | --- | --- | --- |
| ALS2^1^ | Acetolactate synthase | CTTTGGCTCATGGAACGA  CTGCCCAACACCTGTGC | CATATTGTGGCTGGATCTCCTCATTAGAT | 154 | 59 |
| RFP^1^ | Red fluorescence  protein | CGATGGCGACTCTTTCATCT  CGTATAATGGCCACCGTTCT | TGCCACCCACCACACTCATAC | 184 | 60 |
| PP2A^2^ | Serine/  threonine  protein  phosphatase | CCGATCTGTGATATGGGACG  CTAGACCCAAAAGCAACCCA | TGGTTGGTTTTTGTGCGTCTGGCCGG | 209 | 60 |

### ^1^ *ALS* and RFP probes: labeled with FAM^TM^, double-quenched with ZEN^TM^ and Iowa Black Hole Quencher^®^

### ^2^ *PP2A*: labeled with HEX^TM^, double-quenched with ZEN^TM^ and Iowa Black Hole Quencher^®^

**Supplementary Table** **7.** Description and references for construct elements used for transformation

| **Symbol** | **Name** | **Reference** |
| --- | --- | --- |
| *Zm-Wus2* | Maize *wuschel2* | (Lowe et al. 2007) |
| *Zm-Bbm* | Maize *baby boom* | (Gordon-Kamm et al. 2005) |
| *Zs-GREEN* | Green fluorescence protein, *Zoanthus* *sp*. | (Matz et al. 1999) |
| *moCRE* | Maize-optimized CRE recombinase | (Odell et al. 1990) |
| *loxP* | Recombinase target site for CRE recombinase | (Odell et al. 1990) |
| *Sb-ALS_pro_* | Sorghum ALS promoter | SB-ALS promoter and 5’UTR, DOE-JGI Sbi v3.1, SBChr04, bases 49239164-49240031. |
| *Zm-Axig1_pro_* | Maize auxin-inducible promoter 1 | (Garnaat, Lowe, and Roth 2005), NCBI accession AR883375.1 bases 1-1240 |
| *Zm-PLTP_pro_* | Maize phospholipid transferase promoter | GenBank sequence MN380778 |
| *Sb-UBI_pro_* | Sorghum ubiquitin promoter | Unpublished Corteva Agriscience sequence |
| *Zm-Glb1 _pro_* | Maize globulin 1 promoter | (Liu et al. 1998) |
| *Zm-Hsp26_pro_* | Maize heat shock promoter | Unpublished Corteva Agriscience sequence |
